# Sensitivity of Non-Invasive Motor-Unit-Based Gesture Recognition to Signal Degradation

**DOI:** 10.1101/2025.09.10.675419

**Authors:** Mansour Taleshi, Dennis Yeung, Francesco Negro, Ivan Vujaklija

**Author notes:** Corresponding Author: Ivan Vujaklija. The authors are grateful to the Aalto Science-IT project for providing computational resources. The cluster computing tools supplied by this project were instrumental in facilitating the motor unit decomposition and analysis for our study.

## Abstract

The information encoded by motor units has been successfully harnessed to establish high-fidelity human-machine interfaces (HMI). However, the sensitivity of these interfaces using high-density surface electromyography decomposition, a prevalent method for observing motor unit behaviour, to signal degradations commonly encountered in practical settings remains unexplored. Here, we investigated the effects of additive white Gaussian noise (WGN), channel loss, and electrode shift on pseudo-real-time MU-driven motion classification. Across six wrist movements by 13 participants, we evaluated the performance of two classifiers: linear discriminant analysis (LDA) and deep neural networks (DNN), under different noise conditions. The results indicate that spatial perturbations, including channel loss and electrode shift, significantly affected classification accuracy, with LDA being more susceptible than DNN. Conversely, under intense signal noise (WGN with 5 dB SNR), LDA outperformed DNN, and its simplicity potentially provides greater robustness in a challenging environment. Hence, application-specific signal processing considerations are required depending on the target HMI application.

**IMPACT STATEMENT:** Perturbation severely impairs non-invasive motor-unit-based gesture recognition. High signal fidelity and robust system design are essential for practical human-machine interaction.

## I INTRODUCTION

**G**ESTURE-based human–machine interaction (HMI) is reshaping how we interact with Internet-of-Things systems, immersive rehabilitation protocols, and powers intuitive wearable interfaces [1]. Among the diverse biosignals explored for gesture decoding including inertial and optical measurements, surface electromyography (sEMG) provides a direct and temporally precise measure of neuro-muscular activity [2]. These characteristics make sEMG an effective means for real-time gesture decoding, especially in environments where optical methods may be limited [3]. For applications that demand proportional, multi-degree-of-freedom control such as dexterous prosthetic hands and robotic exoskeletons, the field has transitioned toward utilizing high-density sEMG (HD-sEMG), whose rich spatial sampling captures the full spectrum of muscle activity [4].

HD-sEMG arrays record tens to hundreds of channels around the forearm and provides the resolution necessary for blind-source-separation (BSS) algorithms to extract individual motor-unit (MU) spike trains non-invasively [5]–[7]. A common strategy for MU-based HMI application employs a two-stage process for MU decomposition (MUD), where offline/batch MUD stage examines the full dataset to derive MUSTs and generate personalized subject filters and parameters [7], [8], which often referred to as the calibration stage. These parameters are subsequently utilized by a lightweight online MUD that processes incoming EMGs in short time-windows for near-instant MU derivation [8]–[11].

Similar to traditional EMG interfacing, the decomposition of EMG signals can be affected by noise during signal collection. These disturbances may stem from environmental factors, instrument malfunctions, or human-induced errors. Specifically, the dynamic nature of human movements and daily changes in anatomical geometry can lead to electrode omissions (channel loss) and relative motion between the electrode and the underlying skin/muscle [12]– [15]. Additionally, electrode impedance can change due to perspiration or degradation of conductive gel properties affecting EMG signal amplitude and frequency (signal noise) [16]. Although advanced signal processing methods address these issues [17], they often compromise between noise reduction and neural information preservation [18].

This paper examines the impact of spatial and signal-imposed noise on pseudo-real-time motion classification using features derived from a convolutive BSS-based MU-decomposition algorithm. Specifically, we examine the impact of additive white Gaussian noise (WGN), channel loss, and electrode shift on the classification accuracy of linear discriminant analysis (LDA) and deep neural network (DNN) classifiers. By simulating real-world signal degradations, we aim to identify how these non-ideal conditions affect classifier performance and provide insights into optimizing real-time MU-decomposition-based systems for practical applications.

## II RESULTS

To quantify how common recording perturbations compromise MU–based gesture decoding, we recorded 192-channel HD-sEMG from the forearms of thirteen participants while they executed three repetitions of six wrist motions. Motor units were decomposed offline to obtain subject-specific filters, which were then applied online in sliding windows to mimic real-time conditions. Two classifiers were trained on the noise-free baseline (clean) calibration data, including a conventional LDA model and a DNN that uses the richer MU feature space, and were subsequently tested under various perturbed conditions.

Figure 2 displays the classification accuracy (CA) of classifiers for all subjects across all perturbation conditions. The perturbation conditions include additive WGN with various intensity scale (5, 10, 15 dB) and type (constant, linearly increasing, and localized), 15% channel loss, and medial/lateral electrode shift by 1cm. Noise introduction consistently reduced CA across all classifiers and noise types. In clean condition, the DNN classifier achieved a mean accuracy of 89.17% ± 3.21% (95% CI) significantly outperforming (*p* < 0.001) LDA (77.76% ± 3.92%), with an improvement of 11.41%. Under constant WGN at 5 dB, the mean CA for LDA dropped significantly (p < 0.001) to 38.32% ± 8.12%, and for DNN to 27.95% ± 5.65%. With constant WGN at 10 dB, the mean CA decreased to 60.10% ± 5.74% for LDA and to 57.25% ± 7.12% for DNN, both still significantly lower than their respective performances in the clean condition (*p* < 0.001). At 15 dB, LDA achieved a mean CA of 73.66% ± 4.38%, and DNN reached 81.63% ± 5.11%, showing improved performance with higher signal-to-noise ratio (SNR) levels but still below the clean condition.

**Fig. 1:**
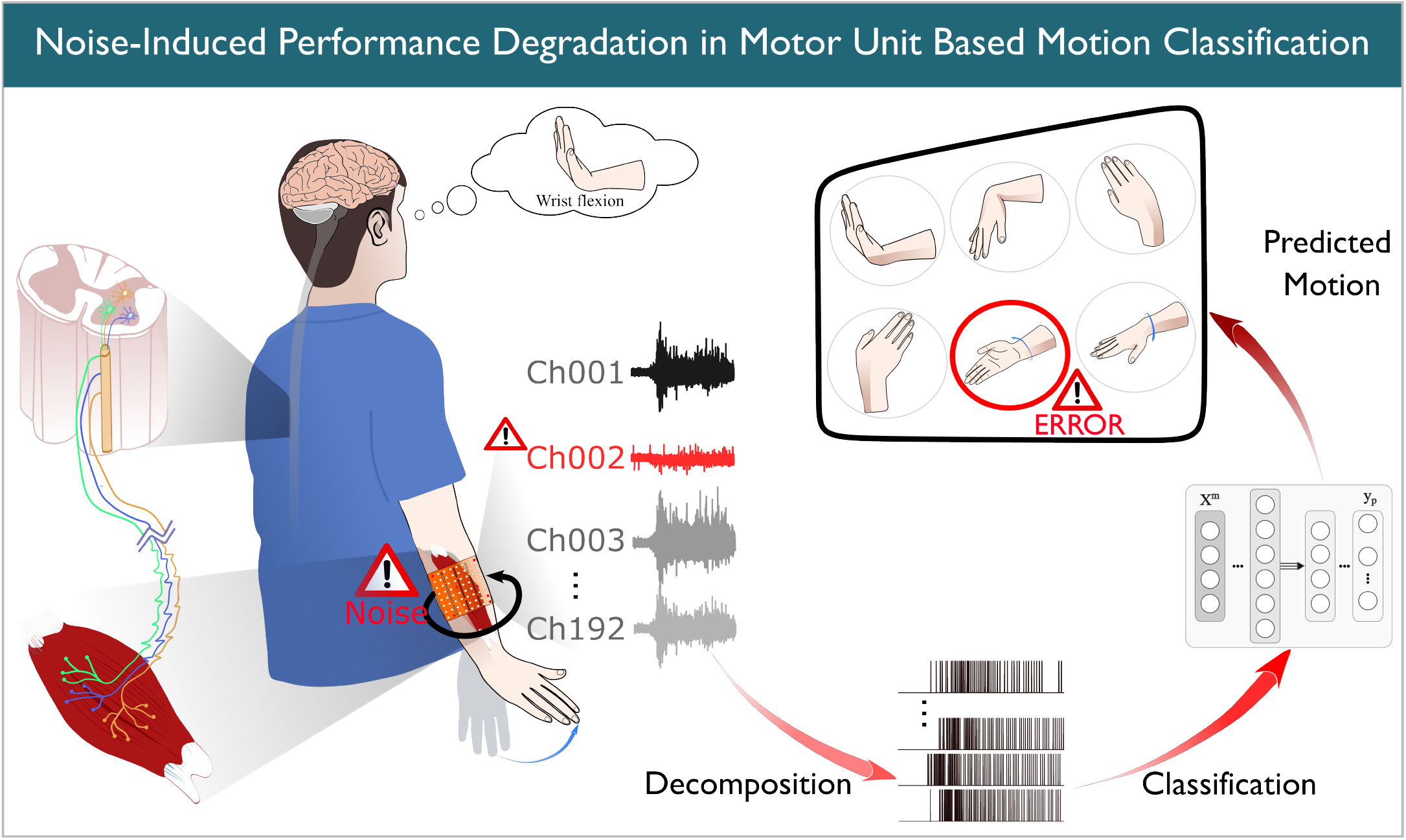
Visual summary illustrates how noise degrades the accuracy of motor-unit-based motion classification. High-density surface EMG recordings (Ch001 to Ch192) capture the activity of individual motor units in forearm muscles during an intended wrist flexion task, but the introduction of noise into channels distorts the raw signals. After decomposition, the extracted motor unit spike trains are passed to a machine-learning classifier that maps firing patterns to predicted movements. The classifier is susceptible to erroneous motion outputs when noise perturbs the input signals.

**Fig. 2:**
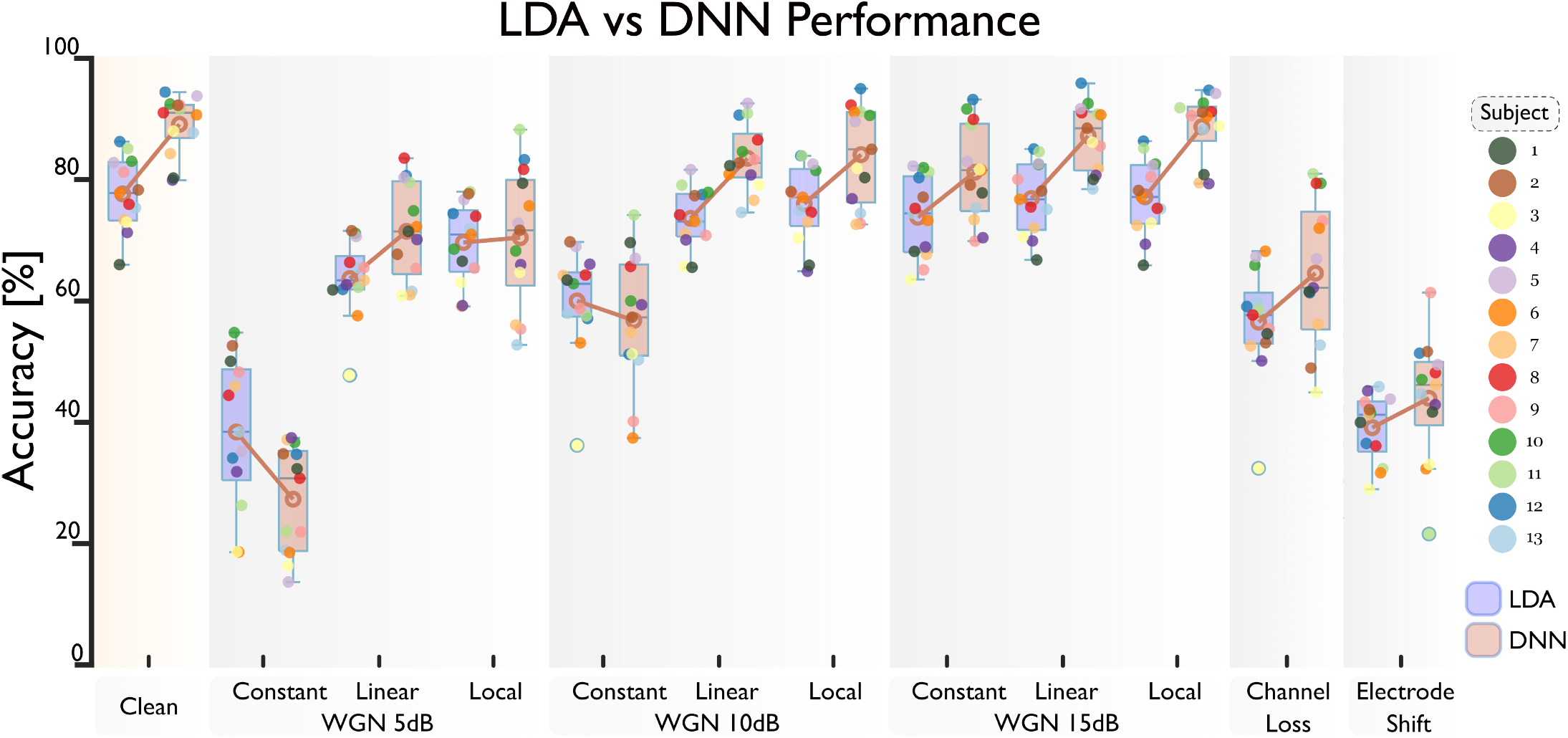
Classification accuracy across all perturbations is shown for each of the 13 subjects via LDA and DNN models under clean conditions, WGN at 5 dB, 10 dB and 15 dB (with constant, linear and local noise conditions), channel loss, and electrode shift. Dots indicate individual subject CA scores, brown lines show the mean CA. LDA, DNN and WGN stand for linear discriminant analysis, deep neural network and white Gaussian noise, respectively.

For linearly increasing WGN at 5 dB SNR, LDA’s mean CA decreased significantly (*p* < 0.001) to 63.46% ± 4.18%, and DNN’s to 72.23% ± 4.90%. At 10 dB SNR, the mean CA improved to 73.59% ± 3.31% for LDA and 84.17% ± 3.18% for DNN, though still slightly below the clean condition. At 15 dB SNR, LDA achieved 76.96% ± 3.91%, while the DNN reached 87.86% ± 3.24%, which is close to, yet does not exceed, the clean-condition performance.

Under localized WGN at 5 dB SNR, LDA maintained a mean CA of 69.39% ± 4.36%, and DNN achieved 71.84% ± 6.61%. At 10 dB and 15 dB SNR levels, both classifiers showed improved performance, with LDA achieving mean CAs of 76.05% ± 4.18% and 77.26% ± 4.06%, with DNN achieving mean CAs of 84.85% ± 5.01% and 88.68% ± 3.56%, respectively, which are closer to the clean condition but still indicate a slight decrease. Channel loss (15% channels lost) significantly decreased CA compared to the clean condition, dropping to 56.16% ± 6.07% for LDA and 65.50% ± 7.56% for DNN (*p* < 0.001). Similarly, electrode shift (1 cm displacement) resulted in significant reductions, with LDA at 38.48% ± 3.39% and DNN at 43.82% ± 6.75% (*p* < 0.001).

The ability of both classifiers to predict specific motions across all conditions are shown in Figure 3 using the F1-score as a metric. Wrist radial deviation and forearm supination and pronation were more challenging for both classifiers. In constant 5- and 10-dB WGN conditions, the LDA classifier generally outperformed the DNN across most motions.

**Fig. 3:**
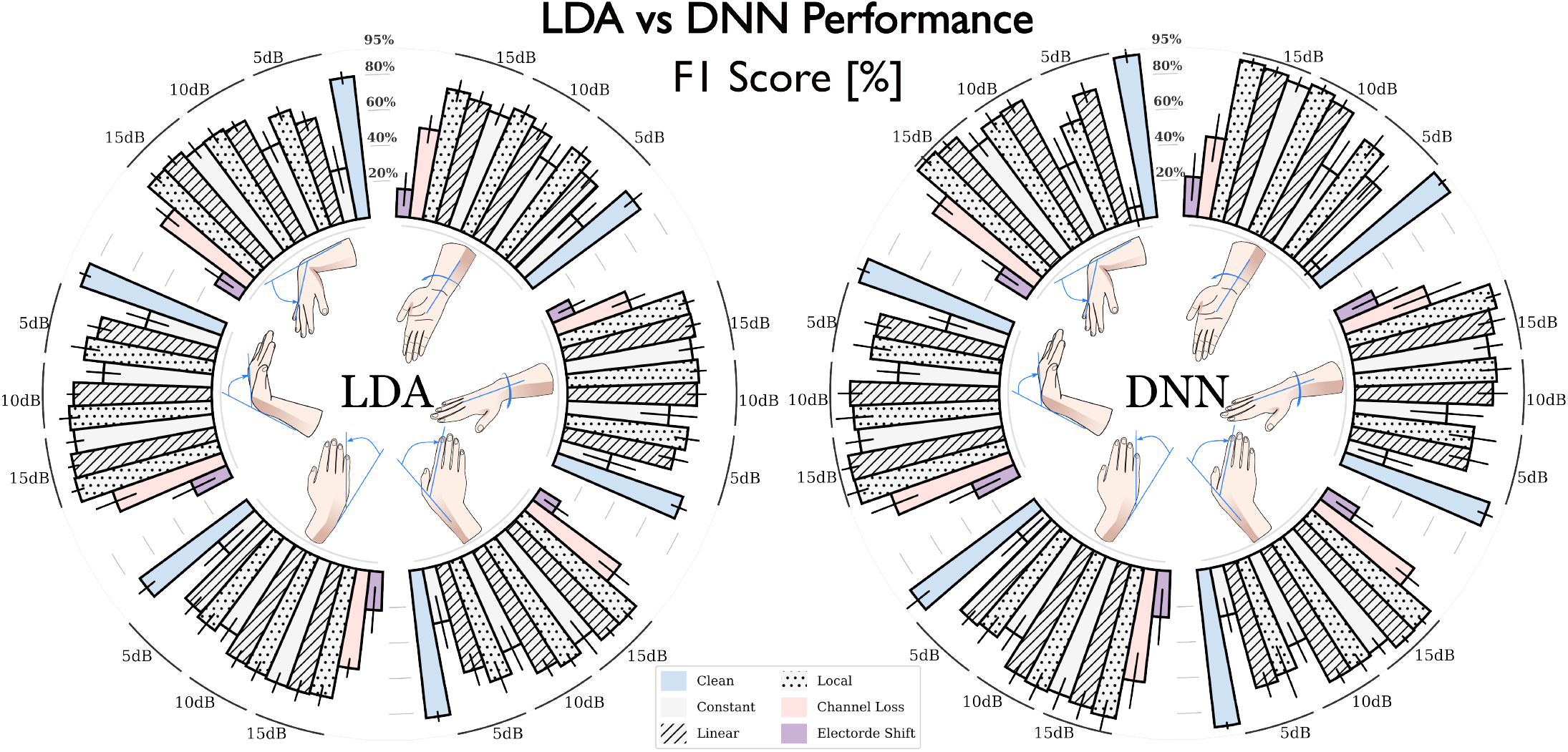
F1 score (%) across 3 noise types (11 variations) using linear discriminant analysis (LDA) and deep neural network (DNN) classifiers. The six motions, shown around the inner circle, are arranged anticlockwise from the top (12 o’clock): wrist flexion, wrist extension, wrist radial deviation, wrist ulnar deviation, forearm pronation, and forearm supination.

## III DISCUSSION

This study quantified the sensitivity of MU-based gesture recognition to common real-world noise conditions simulated through additive WGN, channel loss, and electrode shift perturbations. We specifically analyzed the impact these simulated conditions have on LDA and DNN MU-driven-classifier performance. The results indicate that spatial perturbations, particularly electrode shifts and channel loss, significantly degrade classifier performance, while the effect of signal-based noise depends on its amplitude and distribution, with higher noise levels causing more substantial decreases in CA.

Under the clean condition, the DNN classifier outperformed the LDA, achieving a mean CA of 89.17% compared to 77.76%. This suggests that DNNs can effectively capture complex patterns in MU features extracted from HD-sEMG signals. However, when exposed to certain perturbations, such as 5 dB WGN and 10 dB constant WGN, the LDA maintained higher CA than DNN. This indicates that simpler classifiers may be less sensitive to some forms of noise.

### A. Batch and Pseudo-Real-Time Motor Unit Extraction

Disturbances such as 5db WGN (all types), 10db WGN (constant), channel loss, and electrode shift, can significantly alter feature representation, and hinder LDA and DNN from effectively separating the data. The impact of local and constant WGN on CA is consistent with findings in [13], [19], which report a 10% reduction from a single noisy channel, with performance degrading further as more channels are affected. The channel loss perturbation findings align with Dai et al. [20], who showed a decrease in CA as the number of EMG channels was reduced. Additionally, [19] found that electrode shifts in any direction degrade CA in classifiers using time-domain features.

An alternative interpretation is that the features derived from MU decomposition are inherently sensitive to spatial perturbations due to the localized nature of MU action potentials. The fact that both classifiers suffered under electrode shift conditions suggests that the degradation is not solely due to the classifiers themselves but also to the loss of critical spatial information in the MU features. Generally, during the testing of the motion classifier, the addition of different perturbations to the EMG matrix resulted in the failure of the pre-calibrated whitening matrix *W* and cluster centroids, all developed under clean conditions, in producing accurate observations and threshold for MU detection in noisy environments, causing a significant decrease in CA depending on the noise level and type. An adaptive rule for updating these parameters could potentially address this issue [21].

Our findings emphasize the challenges of deploying MU decomposition algorithms outside laboratory settings. The significant sensitivity to spatial perturbations indicates that ensuring stable electrode placement is crucial for reliable performance. Moreover, the results suggest that while advanced classifiers like DNNs hold promise, their implementation must consider environmental robustness. A major limitation of the present work is that the perturbations were simulated, well defined, and in isolation. Real-world use can involve mixtures of noise sources, time-varying electrode-skin impedance, and larger or irregular displacements that may interact in ways not captured here, at the same time target user populations (e.g., individuals with motor impairments) may differ substantially from the cohort studied here. We also compared only two classifiers-LDA and a GRU- and other architectures (for example, convolutional, or graph-based networks) might exhibit different robustness profiles. Additionally, the simulated perturbations applied to able-bodied volunteers without user-in-the-loop adaptation.

Addressing these limitations will require testing in unconstrained environments with diverse model architectures, integrating adaptive learning from user feed-back [10], [21], and incorporating simulated or real-world/psychological perturbations during training to improve robustness in real-world applications [22]. Such advances should help bridge the gap between the high accuracies achievable under laboratory conditions and the reliability demanded for everyday human–machine interfacing.

## IV CONCLUSION

This work assessed how three varied-scale EMG signal disturbances yielding 11 perturbation conditions affect motor-unit-based motion classification using HD-sEMG data from 13 participants, assessed with both an LDA and DNN. We observed that spatial-based noises, including channel loss and electrode shift, highly affected the classification performance. In contrast, the effect of signal-based noises depends on their amplitude and globality. We have also found that while DNNs exhibit higher accuracy under clean conditions, LDA was more immune to certain noisy scenarios. These results highlight that strong lab performance does not guarantee real-world reliability. Practical MU-based systems will require stable electrode placement, ongoing channel quality monitoring, and adaptive methods to manage signal variability. Future work should explore dynamic noise conditions, broader model types, and validation with users who have neuromuscular impairments.

## V MATERIALS AND METHODS

### A. Participants

Thirteen able-bodied subjects with no known neurological or musculoskeletal disorders participated in this study. The study cohort was comprised of four females and nine males (12 right-handed, one left-handed), aged 33 years on average (range 29 − 37 years). The Ethics Committee of Aalto University granted ethical approval for this study, and all participants signed an informed consent form before committing to the experiment.

### B. Signal Acquisition and Experiment Setup

Joint kinematic and HD-sEMG signals from each subject’s dominant arm were collected. HD-sEMG was recorded using three 8 × 8 (192 channels, each 4*mm* diameter with 10*mm* inter-electrode distance) electrode arrays (ELSCH064NM3, OT Bioelettronica, IT) that covered the upper half of the forearm. This arrangement allowed for monitoring the majority of muscles involved in all degrees of freedom (DOF) of the wrist (Figure 4). A ground electrode and the reference electrodes for arrays were placed on the right wrist. EMGs, sampled at 2048*Hz* and digitized at 16 − *bit* by a Quattrocento bio-amplifier (OT Bioelettronica, IT), underwent in-hardware band-pass filtering (third-order Butterworth, 3 − 900*Hz* cut-off frequency). Alongside, joint kinematics were recorded by three wireless inertial measurement units (IMUs) (MTw Awinda, Xsens Technologies B.V, NL) on the upper arm, lower forearm, and hand, synchronized with the HD-sEMG system at 80*Hz* (Figure 4).

**Fig. 4:**
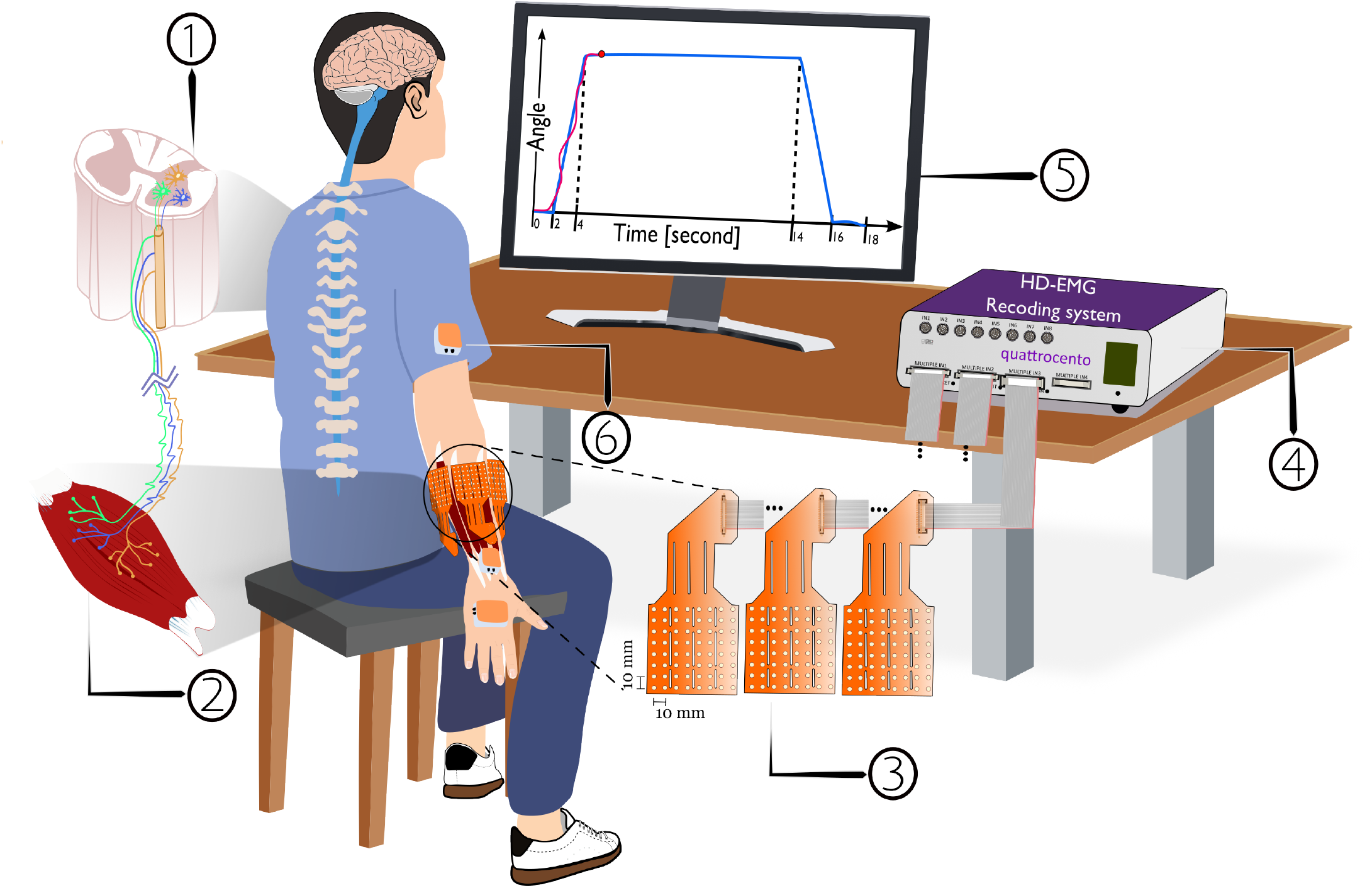
Muscle fibers are physically innervated by alpha motor neurons (MN) of the spinal cord, and their contraction pattern is regulated by this synaptic connection (1,2). The contraction of skeletal muscle fibers can generate electrical activities which can be recorded as EMG signals (3). Due to the one-to-one relationship between action potential triggered in the alpha MN and action potentials triggered in the innervated fibers, EMGs contain information about alpha MN activities and are considered as a function of the total neural input to the muscle. Here, three 8 × 8 electrode arrays are placed around the circumference of the wrist to record the electrical activities from most muscles involved in wrist movements (3). Each electrode has a diameter of 4*mm* and was arranged at a 10*mm* inter-electrode distance, both horizontally and vertically (3). Electrodes are also connected to the multi-channel amplifier and, alongside joint motion, are recorded (4). Real-time feedback of joint motion and desired trajectory is visually provided to subjects in the monitor (5). Joint kinematics were recorded by three inertial measurement units (IMUs) on the upper arm, lower forearm, and hand (6).

Moreover, additional measures were taken to ensure signal quality and maximize the signal-to-noise ratio. Data collection occurred in a magnetically shielded room with a separate electric grid that minimized power supply noise and cross-contamination. Participant forearms were shaved as needed and had their skin cleaned with alcohol. Due to small forearms or poor electrode contact, on average 15 out of 192 channels (range 0 − 42 channels) were excluded from the raw data following visual inspection.

Participants performed three repetitions of three DOF motor tasks: wrist flexion/extension, wrist radial/ulnar deviation, and forearm pronation/supination, chosen for their prevalence in daily tasks [19], [23]. A trapezoidal contraction cue visually prompted each motion, with two seconds of rest, two seconds of ramping, and ten seconds of steady contraction at a comfortable level for the full range of the respective DOF. Subjects viewed the desired and actual trajectory cues on a screen and received real-time feedback on joint motion relative to the desired trajectory (Figure 4).

Both batch and pseudo-real-time MU-decomposition were performed using BSS method [7]–[9], and the process is shown in Figure 5. C and .D.

**Fig. 5:**
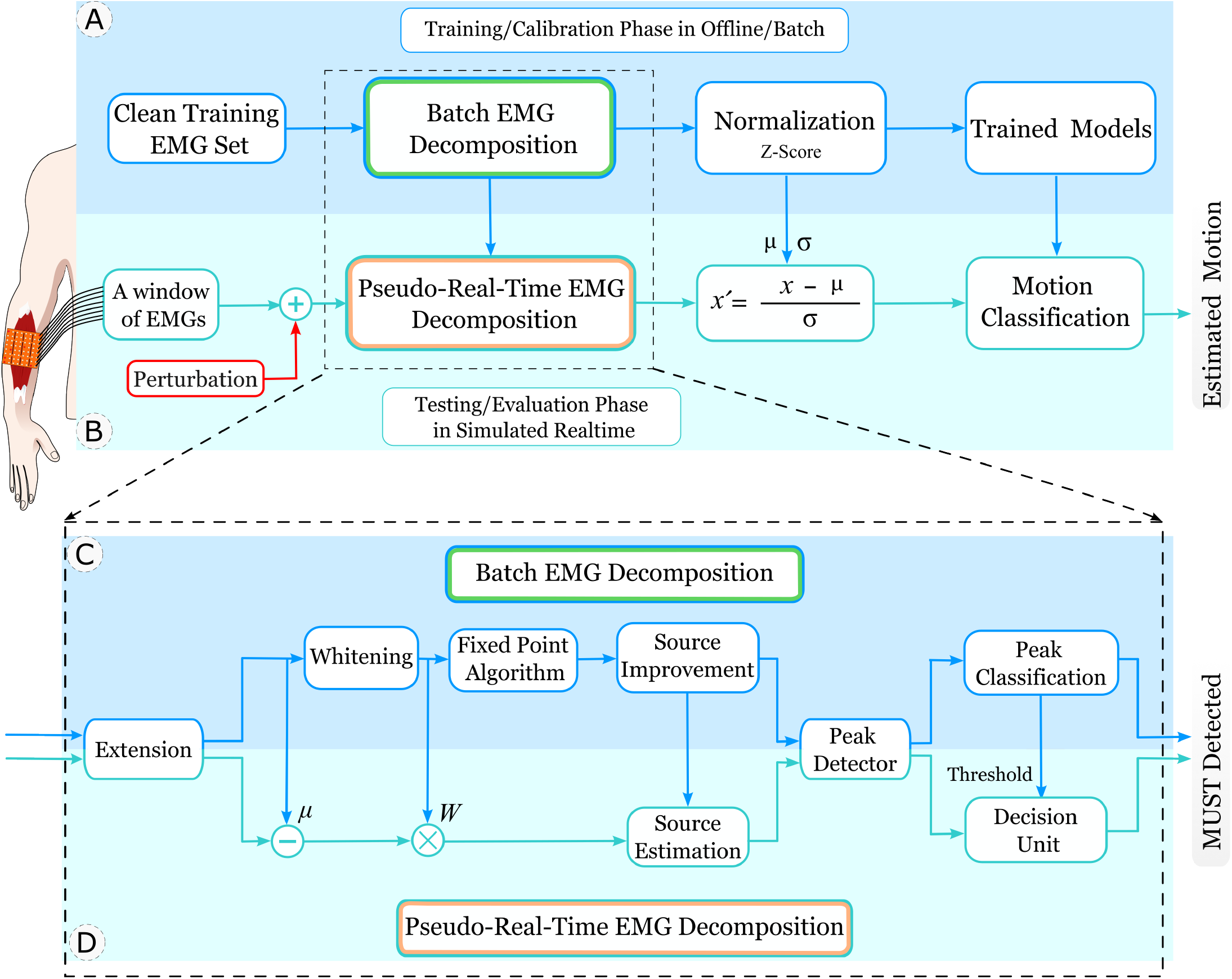
Motor-unit-based motion classification pipeline. The top panel (A) shows the training/calibration phase in the offline/batch manner. Motor unit spike trains (MUSTs) are extracted from the clean training set, then normalized using the Z-score normalization method, and fed to classifiers. The top panel (B) illustrates the testing/evaluation phase in a simulated real-time manner. Perturbations are added to a window of EMGs, then MUSTs are extracted using information obtained in the calibration clean phase. Extracted MUSTs are normalized using the mean (*µ*) and standard deviation (σ) of the training features, and applied to the trained model to estimate motion. The bottom panel (C and D) shows MUSTs extraction pipeline. Batch EMG decomposition (C) performs the computationally demanding tasks of MUST extraction. W is the whitening matrix (from batch SVD) applied to mean-subtracted, extended signals prior to source estimation. *µ* is the channel-wise mean vector. For real-time applications, this step establishes system parameters required by the pseudo-real-time phase (D) to minimize computations of MUSTs extraction.

During the batch decomposition the multi-channel EMG signals were extended by a factor *E* = 10 [11], [24], then whitened via singular value decomposition (SVD) [6], crucial for transforming signals into uncorrelated, unit variance sets. Next, sources were estimated by maximizing non-Gaussianity and sparsity, using fixed-point algorithms with Gram-Schmidt orthogonalization to ensure uniqueness [7]. MUSTs are estimated from squared sources using peak detection and K-means clustering, forming spike train representations [7].

Decomposition quality was verified by silhouette (SIL) measure and coefficient of variation (CoV) for the interspike interval (ISI), accepting only MUs with *SIL >* 0.9 and *CoV*_*ISI*_ *<* 30% [11]. SIL, a normalized EMG decomposition accuracy indicator, differentiates intra- and inter-cluster point-to-centroid distances. This method, validated with experimental and simulated signals, involves a final manual inspection of MUs to ensure reliable discharge patterns for further analysis [6], [7]. The pseudo-real-time decomposition, suitable for real-time HD-sEMG signal processing, utilizes parameters from the batch phase to guarantee a certain processing time [9], [11], [25], [26]. Processing in 160*ms* windows without overlap, it involves extending signals, subtracting the mean, applying the whitening matrix, and using matched filter banks for source signal estimation [9], [11], [26]. Detected peaks from squared sources signals are classified as MUST or noise based on proximity to cluster centroid [9].

### C. Perturbation Conditions

In practical HD-sEMG deployments, electrode arrays and acquisition systems are exposed to a variety of unpredictable disturbances that degrade signal quality and downstream MU decomposition. To mimic these real-world challenges, we defined eleven perturbation scenarios drawn from three broad categories and applied them to the raw EMG recordings before feeding MU features into the classifiers. Since practical noise sources (e.g., cable interference, amplifier thermal noise, impedance fluctuations) are difficult to predict in advance, our simulations cover a variety of noise intensities and spatial distributions to systematically evaluate the sensitivity of MU-based classifier.

#### C.1. White Gaussian Noise (WGN)

Additive WGN approximates a combination of thermal noise in the electronics and time-varying impedance changes at the electrode–skin interface (caused by gel drying, sweat, or micro-motion) that collectively raise the noise floor and reduce SNR over a recording session [13], [16], [27], [28]. To capture both global and localized noise effects, we contaminated EMG recording with three WGN variants: 1) a constant WGN on all EMG channels, 2) a progressively increasing disturbance on all EMG channels, and 3) a constant WGN on a local 4 × 4 cluster of channels with a randomized location. Each variant was tested at three SNR settings (5*dB*, 10*dB*, and 15*dB*), which resulted in nine distinct noise profiles. WGN is termed ‘signal-based’ as it alters raw signal attributes without affecting sensor placement or losing sensor data.

#### C.2. Channel Loss

To address the challenge of unusable channels in HD-sEMG, often due to poor electrode-skin contact, disconnection, or low-amplitude signal detection issues [13], [14], we simulated this by setting about 15% of electrodes (29 out of 192) to zero during the classifier’s testing phase. This reflects real-world scenarios where classification performance is impacted by the loss of critical information from these channels.

#### C.3. Electrode Shift

Electrode displacement or shift is another frequently noted perturbation in surface EMG signals [13], [15]. Such displacement, often caused by human activities, significantly reduces model performance and impacts both industrial and real-world prosthetic applications. This issue is a key factor for the discrepancy in performance between real-world and controlled laboratory settings, where myographic control pattern recognition has seen considerable advancements. To simulate this, we rotated all electrodes in the 8 × 8 arrays by one column, equating to an eight-channel shift or a 10*mm/*1*cm* medial/lateral displacement, considering the inter-electrode distance.

### D. Motion Classification under Perturbation Conditions

Here LDA and DNN classifiers were trained during the training/calibration phase using batch MUs features. The performance of the trained model was then tested in a simulated real-time environment with EMG perturbations using pseudo-real-time MUs features (Figure 5). In test phase, pseudo-real-time MUs features were extracted based on parameters from batch decomposition.

Extracted features underwent z-score normalization before classification. A 3-fold cross-validation assessed the estimator’s prediction accuracy on new data, alternating between two repetitions for training and one for testing. To evaluate LDA and DNN classifiers, CA and the F1-score, a normalized harmonic mean of precision and recall, were used to measure overall quality and class-specific performance, respectively. Our 9-layer deep Gated Recurrent Unit (GRU) architecture, part of the DNN classifier, includes sequential, GRU, leaky ReLu, dropout, GRU, leaky ReLu, fully connected, softmax, and classification layers.

### E. Statistical Analysis

In the study, statistical analyses assessed significant differences in pseudo-real-time MU-decomposition driven motion classification under perturbations. The Kolgomorov-Smirnov test was used to check data normality. Pairwise comparisons between conditions were conducted using paired t-tests. To control for multiple comparisons, p-values were adjusted using the Bonferroni correction. The significance level was set at *p* = 0.001.

